# Multi-Link Analysis: Brain Network Comparison via Sparse Connectivity Analysis

**DOI:** 10.1101/277046

**Authors:** Alessandro Crimi, Luca Giancardo, Fabio Sambataro, Alessandro Gozzi, Vittorio Murino, Diego Sona, the Alzheimer’s Disease Neuroimaging Initiative

## Abstract

The analysis of the brain from a connectivity perspective is unveiling novel insights into brain structure and function. Discovery is, however, hindered by the lack of prior knowledge used to make hypotheses. On the other hand, exploratory data analysis is made complex by the high dimensionality of data. Indeed, in order to assess the effect of pathological states on brain networks, neuroscientists are often required to evaluate experimental effects in case-control studies, with hundreds of thousand connections.

In this paper, we propose an approach to identify the multivariate relationships in brain connections that characterise two distinct groups, hence permitting the investigators to immediately discover sub-networks that contain information about the differences between experimental groups. In particular, we are interested in data discovery related to connectomics, where the connections that characterize differences between two groups of subjects are found. Nevertheless, those connections not necessarily maximize accuracy in classification since this does not guarantee reliable interpretation of specific differences between groups. In practice, our method exploits recent machine learning techniques employing sparsity to deal with weighted networks describing the whole-brain macro connectivity. We evaluated our technique on functional and structural connectomes from human and mice brain data. In our experiments, we automatically identified disease-relevant connections in datasets with supervised and unsupervised anatomy-driven parcellation approaches, and by using high-dimensional datasets.

## Introduction

The analysis of brain networks, or connectomes, is a recent and exciting advancement in magnetic resonance imaging (MRI) which promises to identify new phenotypes for healthy, diseased or ageing brains^1^. A connectome is a comprehensive map of the connection in the brain, which is conceived as a network, where brain areas (nodes) are connected by links (edges)^2^, and connections can be either given by white matter tracts between pairs of brain regions, or by an index of correlation of functional activity^3^. This allows for analysing the brain as a complex system of dynamically interacting components without explicitly relying on local activation or brain morphology.

### Case-control studies and connectomics

Experiments with connectomes are typically designed by comparing a studied group with a control group in order to identify brain-network topological biomarkers relevant to the studied group^4^. Indeed, inter-group differences in some of these topological measures have been discovered for various neuropsychiatric disorders^5^, like Alzheimer’s disease^6^, multiple sclerosis^7^, schizophrenia^8^, stroke^9^, major depression^10^, autism spectrum disorder^11^, etc. All these approaches use topological measures with statistical tests to assess their discrimination power in a univariate analysis framework. Alternatively, in a multi-variate framework, machine learning methods have been proposed to differentiate groups of subjects using topological measures^12^. Surveys on graph-topological metrics using functional magnetic resonance imaging (fMRI) data and related clinical applications using structural features are given respectively in Varoquaux et al.^13^ and Griffa et al.^14^.

### Local differences between connectomes

The main drawback of the aforementioned approaches is the limited interpretability of graph statistics as they miss the local characterization of the groups in terms of differences in the connectivity, but rather employ global statistics which are difficult to be translated into clinical settings for local analysis. A method which allows insights on local connectivity patterns in case-control studies relies on network-based statistics (NBS). In this approach, the connectivity between pairs of brain regions is tested for significance using univariate statistics for functional^15^ and anatomical^16^ connectivity disturbances. Simpson et al.^17^ extended the NBS method using a permutation test based on Jaccard index at node level. While, Chen et al.^18^ enhanced NBS regulating the topological structures comprised. Other research groups^19–21^ leveraged support vector machines (SVM) weights to identify discriminating regions. SVM is a supervised learning method which constructs a hyperplane or set of hyperplanes in a high-or infinite-dimensional space used for classification. Specifically, Ng et al.^20^ used a projection of covariance estimates onto a common tangent space to reduce the statistical dependencies between elements. Then, while Mastrovito et al.^19^ employed recursive feature elimination (RFE) to identify connections relevant for the classification, both Goankar et al.^21^ and Ng et al.^20^ found meaningful connections using t-tests on the SVM weights. A more advanced machine learning approach is based on SVM coupled with Riemannian/Grassmannian geometry^22^. Van Heuvel et al.^23^ proposed a sub-graph level analysis for a more specific and localized information, with a specific emphasis on the potential functional importance of highly connected hubs (“rich-clubs”). Although the focus on rich-clubs is insightful, this method could leave out subtle differences between case-control groups which are not present in highly connected hubs. Lastly, despite NBS and its extensions have been shown to outperforms other methods in comprehensive comparisons, the identification of graph sub-networks is a pre-requisite which can limit the detected connections and the t-tests are carried out in a univariate manner^24^. Moreover, the choice of related statistics can influence considerably the results^24^.

### Relation to previous methods

In this context, we are interested in data discovery related to connectomics, where the connections that characterize differences between two groups of subjects are found, and where maximizing accuracy does not guarantee reliable interpretation since similar accuracies can be obtained from distinct sets of features^25^. To overcome the limitations of the univariate approaches, which perform statistical tests on single connections as mentioned in the previous subsection - and in particular to the most commonly used NBS^26^ - we use a multivariate bootstrap-like approach followed by a stability selection step. Therefore, we propose a fully data driven method to identify relevant brain sub-networks in experiments with case-control design which can be used as an hypothesis generation tool for connectomes investigations. Our method has the potential to work equally well with functional and structural MRI data, and no prior knowledge about the type of connectivity is required, only examples of brain connectivity matrices of two groups are needed.

A similar method proposed by McMenamin and Pessoa^27^ implemented a two-layer dimensionality reduction technique based on principal component analysis (PCA), followed by quadratic discriminant analysis to identify clusters with altered connectivity at voxel level. However, when PCA was used for feature selection, the eigenvalues of the covariance matrix were used regardless the prior knowledge on the groups to be discriminated, and in doing so the resulting features may not be those which were really meaningful in terms of discrimination between groups. Conversely, our method directly performs a sparse version of linear discriminant analysis (LDA) that, by design, tries to optimize the feature selection step aiming at discriminating the groups. This allows the proposed method to be more specific in terms of identified discriminating connections. Furthermore, sparse models follow a feature selection agenda to subselect among existing variables, whereas PCA dimensionality reduction follows a feature engineering agenda to generate a set of new variables. A feature selection by sparse model is indeed similar to the RFE used by Mastrovito et al.^19^. However, the stability of RFE approach depends heavily on the type of model used for feature ranking at each iteration, and as shown empirically, using regularized ridge regression jointly to stability selection criteria can provide more stable results in terms of *stability selection* of features, and yields finite sample familywise error control^28,29^. More specifically, the proposed model is based on an ensemble of sparse linear discriminant models allowing to find the networks’ elements (a set of edges) able to consistently distinguish two groups, in the attempt to minimize the subset of selected connectivity features and simultaneously maximize the difference between the groups^30^. Essentially, the system acts as a *filter* removing the elements that are not useful to discriminate between the groups. First by enforcing sparsity at individual level. Then, by performing a second stage of feature filtering across the dataset to assure stability selection. This feature selection process is not inherently specific to connectomes as it can be applied to arbitrary high-dimensional, multivariate datasets. Nevertheless, recent studies showed that sparsity based approach can be particularly useful in graph/connectome analysis as they can highlight significant connections when prior knowledge is missing^24,31–33^.

Other methods have already used sparsity to estimate relevant connections^34–36^. However, these methods did not focus on finding the discriminant connections between groups while performing the sparse selection. They use sparsity to reduce the number of connections regardless on the inter-class discrimination.

### Multi-link Analysis (MLA)

The interpretation of differences in brain networks is not always straightforward given individual variability and the high dimensionality of data^37^. Moreover, the internal structure of the brain connectivity with cross-relationships and dependencies in the feature space (the edges) may prevent a full retrieval of groups’ differences using univariate analysis. Machine learning and dimensionality reduction techniques are designed to solve these issues, and hence these methods are a natural choice for addressing this discrimination task. We propose a two-stage feature selection process. In the first stage a classifier reinforcing sparsity is employed to select discriminant features, iterating over different subsamples of the dataset in a bootstrapping framework. Then, only features which are consistently selected across the iterations are kept according to a stability selection criterion.

An approach simultaneously implementing classification and feature selection in a sparse framework is *sparse logistic regression*, which has been already used to select relevant voxels for decoding fMRI activity patterns^38,39^. Alternatively, in case of Gaussian-distributed data, the well known *linear discriminant analysis* has been extended to the sparse case with the *sparse discriminant analysis* (SDA) model^30,40^. In particular, the method by Clemmensen et al.^30^ implements the elastic net regression with the *ℓ*_1_-norm on the feature weights that indirectly sets the number of selected features.

For all the experiments, the connectivity matrices are vectorized and ordered as rows in a *n* × *p* data-matrix **X**, with *n* being the number of observations and *p* their dimensionality. The corresponding classification of objects is encoded into the *n* × ***K*** indicator matrix **Y**, where each cell **Y**_*ik*_ indicates whether observation *i* belongs to class *k*. The SDA proposed by^30^ then finds the discriminant vectors *β_k_* for each class *k* and the vector of scores ***θ***_*k*_ by the convex optimization given by the following regularized linear discriminant formulation

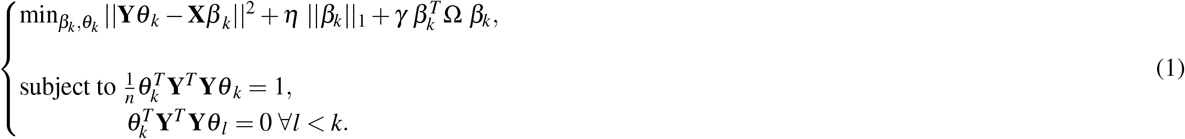

where Ω is an arbitrary positive definite matrix, which allows to calculate a smooth discriminant vectors ***β***_*k*_ even if the number of samples is smaller than the number of features (*n* ≪ *p*). In our experiments we used Ω = **I** which makes the formulation an elastic net problem 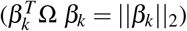. The non-negative parameters ***η*** and ***γ*** control respectively the *ℓ*_1_ and *ℓ*_2_ regularization. The parameter ***η*** can also be reformulated as the number of desired variables which are left in the model, and when used in this context we refer to it as ***α***^41^.

The advantage of the proposed sparse method is its capability of managing high-dimensional data thanks to the *ℓ*_2_ regularization. Moreover, the *ℓ*_1_ regularization term allows the model to select a small subset of features for the linear discrimination. This might result in a loss of predictive power while however reducing the over-fitting problem. In contrast, the *ℓ*_2_ penalty term enjoys the grouping effect property, i.e., it works keeping small and comparable the weights of correlated predictors^41^. Moreover, *ℓ*_2_ penalty term is much better at minimizing the prediction error than *ℓ*_1_ regularization. As a result, their combination allows to determine a good trade-off between an optimal classifier and a minimal selection of relevant predictors. Further details on the regularization parameters are given in the Method section and Appendix.

### Experiments overview

Owing to the sparsity principle driving the learning method combined with the statistical robustness of ensemble methods, our multivariate approach can scale up with the number of analysed connections, even when employing a limited number of whole-brain connectivity matrices. By virtue of being multivariate, this approach can identify brain sub-networks whose edges considered as a set can characterize the differences between the connectomes but taken independently cannot. Moreover, the method does not have to rely on covariance matrices. It just needs an index describing the strength of connectivity between the areas in terms of correlation, similarity, dissimilarity or other metrics. For example, in case of structural connectivity the matrix can be determined counting the number of connections between the areas.

We validated the approach on three real datasets. In a first experiment, we used the structural connectivity, based on the tractography extracted from diffusion tensor imaging (DTI). Specifically, we compared a group of acallosal BTBR mice (a well-characterized model of autism) with a group of control normocallosal and normosocial C57BL/6J mice^42,43^. Performing this experiment with a simple and well known connectivity dysfunction, without the use of any prior anatomical parcellation to avoid any prior bias, we empirically validated the approach, which was able to retrieve the expected dissimilarity between the two groups.

A further experiment was conducted on structural connectivity matrices from a publicly available dataset of patients affected by Azheimer’s disease, where connectivity is also defined by tractography. The final experiment was carried out on a large functional dataset of attention deficit hyperactivity disorder (ADHD) children compared to typically developing (TD) children. Further details are given in the Method section.

In all cases, our method successfully detected inter group differences relevant to the medical condition investigated. Those results are compared to the results obtained by using NBS, and a framework based on SVM weights^20,21^. NBS and MLA select discriminative features in different ways. NBS performs univariate t-tests among the features while MLA performs a sparse multivariate regression. Nevertheless, NBS is the commonly used algorithm for this type of analysis and considered the state-of-art. The SVM based method is a further machine learning approach where we investigated the most significant connections obtained from the SVM discrimination weights similarly to previous studies^20,21^. Selected weights are those larger than the 95-th percentile or smaller than the 5-th percentile of a random weight distribution representing the null hypothesis. The null hypothesis for the SVM weights is obtained by performing 1000 random permutations of the labels of the two groups. In our experiments we used the LibSVM toolbox^44^.

All experiments have been conducted in accordance with relevant guidelines and regulations. The human experiments used publicly available dataset. The Alzheimer experiments have been conducted on data previously acquired by the ADNI initiative according to good clinical practice guidelines, US 21CFR Part 50– Protection of Human Subjects, and Part 56–, acquiring both phone and written consent. The data are from different centers, though the umbrella Institutional Review Board that approved the study and protocol: the University of California, San Francisco. The ADHD experiments have been also conducted on data previously acquired for another study, for which the ethics review board of the New York University have granted the ethical approval and for which informed consent was obtained for each subject. The mice experiments have been conducted in accordance with the Italian law (DL 116, 1992 Ministero della Sanitá, Roma) and the recommendations in the Guide for the Care and Use of Laboratory Animals of the National Institutes of Health. Animal research protocols were also reviewed and consented to by the animal care committee of the Istituto Italiano di Tecnologia (permit 2007–2012). All surgical procedures were performed under anesthesia.

## Results

### Mice Structural Connectivity Data

In order to prove the discriminative power of our approach, we tested its ability to correctly distinguish the structural connectomes of two groups of mice (C57BL/6J and BTBR) characterized by previously described white matter alterations, i.e., the presence/absence of the two major neocortical intra-hemispheric tracts: the corpus callosum and the dorsal hippocampal commissure^45^ as shown in Figure 1. Being the structural alteration in the BTBR mice well known, this dataset is used to validate the proposed method. Indeed, the BTBR mice model represents a ground truth of expected differences between the two groups. Over and above, more than the discrimination between the groups, we are interested in empirically assessing the ability of our approach to correctly identify white matter tracts differences in the two groups.

**Figure 1.**
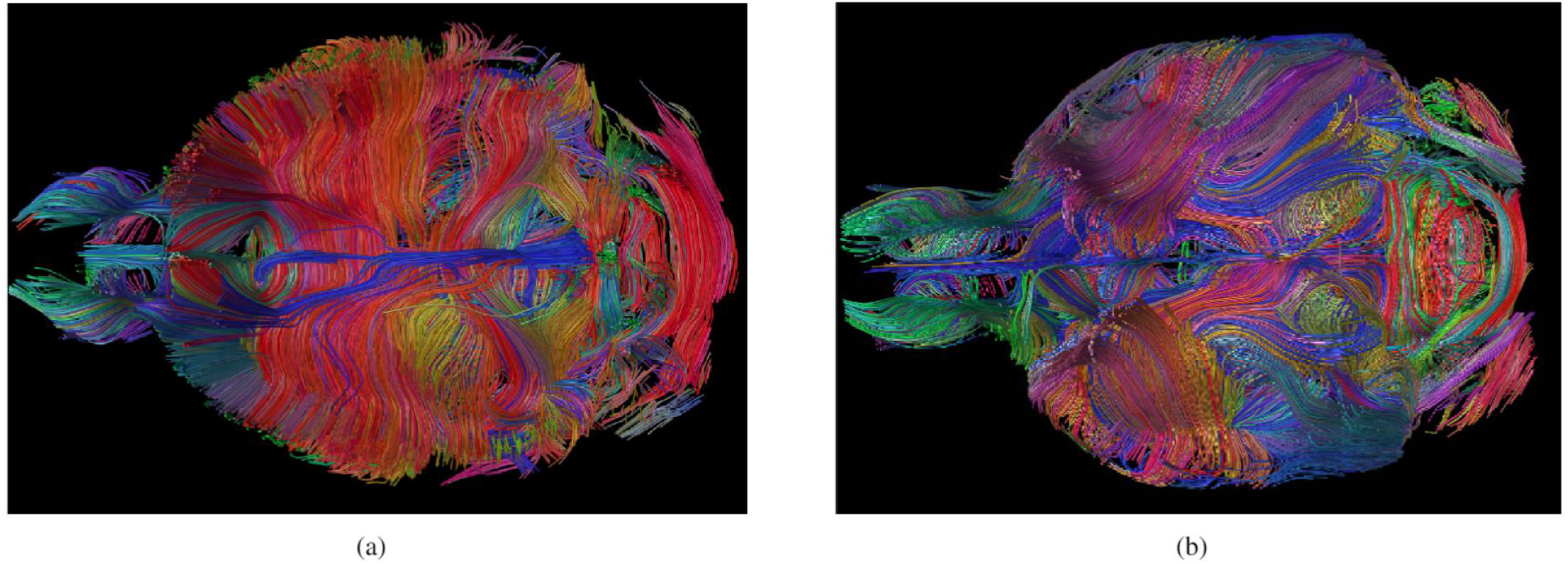
Example of axial section tractography of (a) a normo-callosal C57BL/6J control and acallosal BTBR (b) mouse respectively, where the different anatomical structures are apparent but difficult to understand. In particular, the lack of corpus callous in (b) is visible.

Indeed, by using the proposed algorithm, the model correctly classified all samples in a cross-validation schema, and structural differences - as the lack of corpus callus - were found as expected from literature. The mean misclassification varying the parameter *α* resulting by the cross-validation is shown in Figure 2 (a).

**Figure 2.**
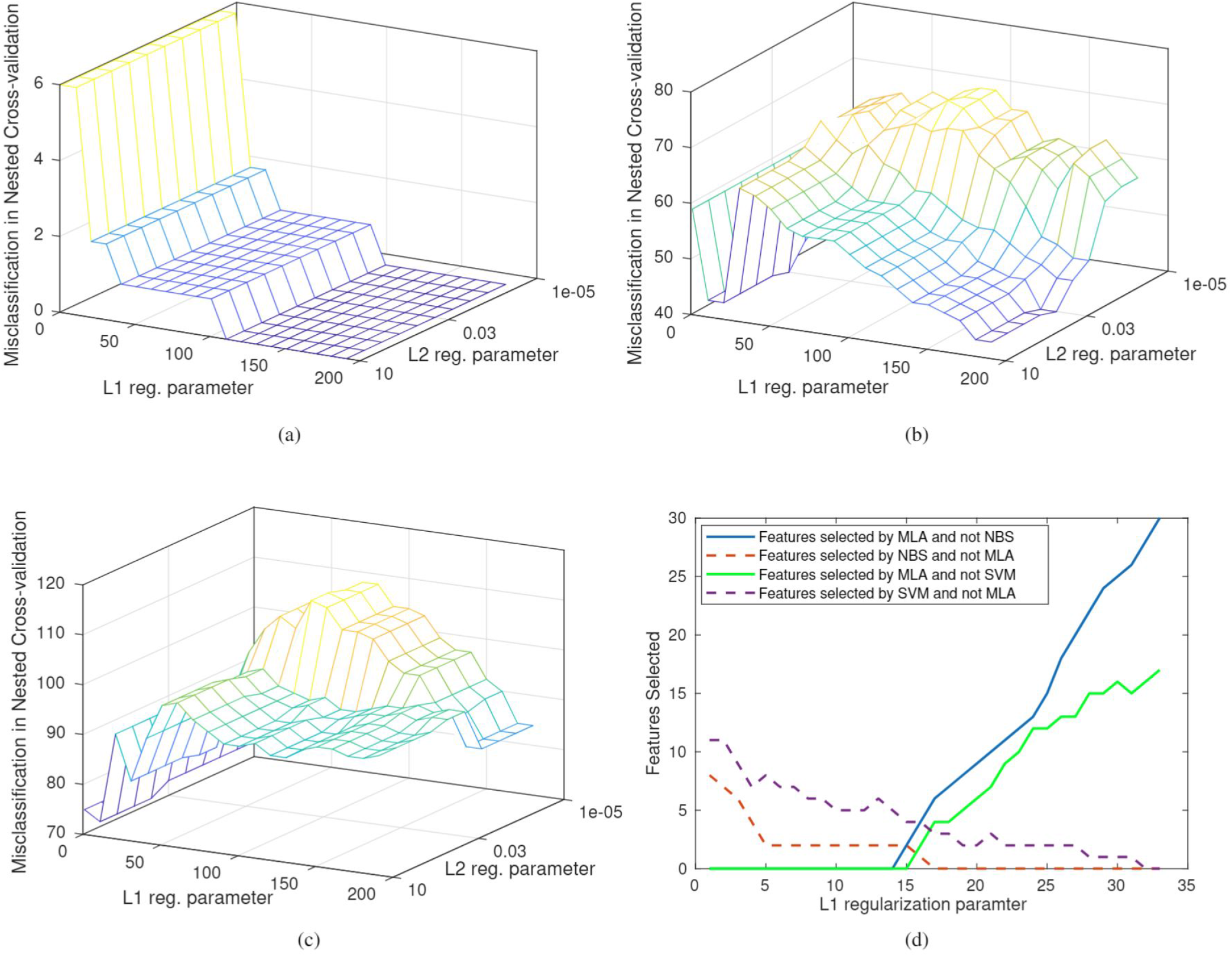
Misclassification error as function of regularization parameter computed with the nested cross-validation. (a) Mice data experiment: The misclassification error reaches a plateau after ***α*** = 110 and the L2 parameter ***γ*** has no influence. (b) Alzheimer human experiment: The misclassification has two plateau, one near ***α*** = 20 and one near ***α*** = 190, and for L2 parameter ***γ*** > 0.03. (c) ADHD human experiment: The misclassification has a plateau near ***α*** = 10, the L2 parameter changes the results but with little influence. (d) Number of features detected by one algorithm and not by the the other varying the amount of sparseness. This graph shows that by decreasing the sparseness, the number of features detected by the MLA is increasing. Example shown for the Alzheimer dataset.

To this aim the proposed approach returns a statistics of the relevance of features, by counting the amount of occurrences of the features selected by the ensemble of models. Figure 3 shows the occurrence of the detected features for the experiment with mice structural connectomes, some of which are present in all the runs, indicating a strong relevance for the problem at hand. Interestingly, the edges identified by the algorithm showed the expected characteristic features of the BTBR strain, including the agenesis of the corpus callosum and the presence of rostral-caudal rearrangement of white matter. Figure 4 shows how our algorithm (MLA) and NBS identify the parts of the corpus callosum which are known to be missing. Results obtained by using the SVM based framework were also similar to those given by NBS. This experiment confirms that our new approach and NBS are able to identify the acallosal connections in the BTBR models.

**Figure 3.**
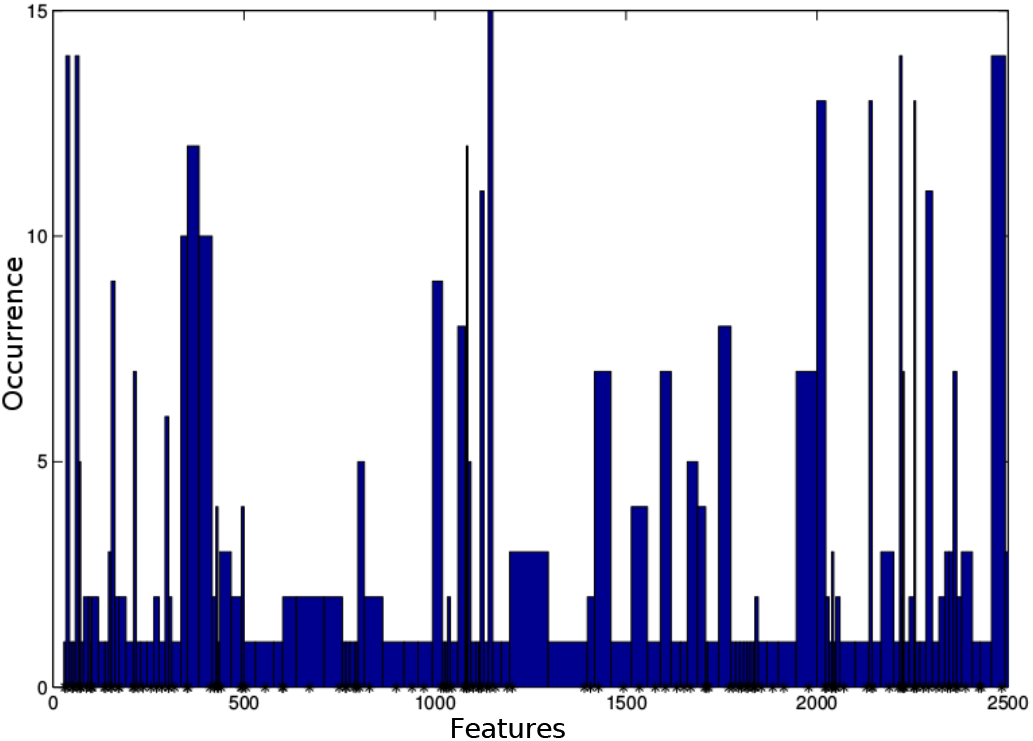
Mice data experiment: Histogram describing the occurrences of features (i.e. brain connections) selected in the mouse experiment. Higher values indicate connections that characterize the differences between BTBR and control mice in the classifiers within our ensemble framework. This information is used to automatically select a sub-set of ”relevant” features. Namely, the most frequent features highlighted by the histogram are kept.

The whole analysis from raw DTI data to tracts selection of the 16 subjects, by using Matlab Mathworks 2014, took less than 40 minutes on a 2.6 GHz machine with 4GB of RAM. However, the five rounds of MLA analysis required only less than 1 sec (with *ℓ*_1_ parameter ***α*** = 110 estimated by nested cross-validation which also gave 100% accuracy).

### Human Alzheimer Structural Data

We tested the algorithm also on a publicly available dataset based on human MRI recorded from patients with Alzheimer against normal elderly subjects. The Alzheimer dataset was used to investigate the influence of the regularization parameter on the set of features selected by the proposed method as compared to NBS (and SVM). In particular, we investigated the ability of the proposed approach to detect features that are not detected by NBS (or SVM) and vice-versa, varying the sparsity parameter ***α***.

The selection of significant features with the NBS algorithm thresholded at p-value = 0.05 produced 14 connections. Using the detected features in a nested cross-validation case-control classification task produced an accuracy of 65%. The selection of significant SVM weights, instead, highlighted 20 connections, which in the nested cross-validation classification gave 66% accuracy. It has to be noted that some of the connections were detected by one algorithm and not by the other.

On the contrary, with a proper choice of the sparsity parameter, the proposed approach detected all features selected by both NBS and SVM and some others. More specifically, as shown in Figure 2 (d), by decreasing the sparsity (i.e., increasing the parameter ***α***), the number of features included by MLA and not by NBS (or SVM) increases, while the number of features detected as relevant by NBS (or SVM) and not by MLA decreases with a break-even-point at ***α*** = 15. The best classification score with the features detected by the MLA was obtained with ***α*** = 33 with an accuracy of 75%. It is worthwhile to mention that generally detected features were symmetric. Namely, if a connection from ROI ***a*** to ROI ***b*** was detected, also the reverse connection from ***b*** to ***a*** was detected. The resulting features produced by MLA approach are depicted in Figure 5. The identified connections were mostly ipsilateral within the two temporal lobes. The analysis of Alzheimer dataset took less than 1 second on a 2.6 GHz machine with 4GB of RAM.

### Human ADHD Functional Data

By using the ADHD dataset, the cross-validation found the optimal solution for the the MLA algorithm at α = 10, highlighting 8 discriminant connections across the groups with an accuracy of 70%. The NBS method, thresholded with a p-value = 0.05, did not find any significantly discriminative connection. The SVM based framework, instead, showed 60% accuracy detecting 2 significant connections.

The inability of NBS to find relevant connections might be due to the fact that its first key step is the identification of candidate subnetworks, which are then tested for their relevance using a permutation test. These candidate subnetworks are selected only when the nodes are well connected each others, however, connectomes determined with high dimensional parcellations (in our case we used an atlas with 200 areas) are more likely to have a sparsely connected network. While being this a problem for methods expecting a densely connected graph, like NBS, it is not affecting our approach that does not have any prior on the expected connectivity. The connections detected by the proposed algorithm are depicted in 6. The ADHD samples analysis took less than 30 seconds on a 2.6 GHz machine with 4GB of RAM.

## Discussion

The proposed method performs a global multivariate analysis characterizing local differences between networks. As this method is based on sparsity principles, it is particularly suited for those experiments with high-dimensional data and small sample size. Moreover, the analysis based on multivariate statistics allows to retrieve sub-networks based on feature dependencies. The limitation of NBS and the SVM-based approach in detecting univariate differences is visible in the experiment with human functional data. In fact, the proposed algorithm detects some connections which are very often selected by the ensemble of learners, as seen in the histogram in Figure 3. On the contrary, with the univariate analysis some edges are discarded as producing non-significant p-values. Nevertheless, MLA, NBS and SVM approaches gave similar results on the experiment with mice data, confirming the starting hypothesis on the anatomical differences between the two mouse lines.

The stability of the selected features is an important characteristic of the algorithm. As assessed empirically, increasing the value of ***α*** the model only introduces new features without dropping any feature determined with smaller ***α***s. The parameter ***α*** represents the strength and limitation of the method. In fact, despite in the reported experiments we determined it automatically through cross-validation, the sensitivity of the algorithm can be manually adjusted through this single parameter which allows the neuroscientists to decide how strong the class characterization should be. A similar user-guided approach with methods based on sparsity has been previously described^34,35^.

When MLA was applied to the acallosal BTBR mice, a mouse model of autism^42,43^, as shown in Figure 4, the tracts detected as discriminant were those with altered white matter connectivity in BTBR mice with respect to control mice. These results are in line with previous results in literature, including the lack of corpus callosum and hippocampal commissure^46–48^, and increased intra-hemispheric ipsilateral connectivity^49,50^, also observed in human patients with autism spectrum disorder (ASD)^51,52^. This demonstrates that the algorithm is able to identify the known differences between groups.

**Figure 4.**
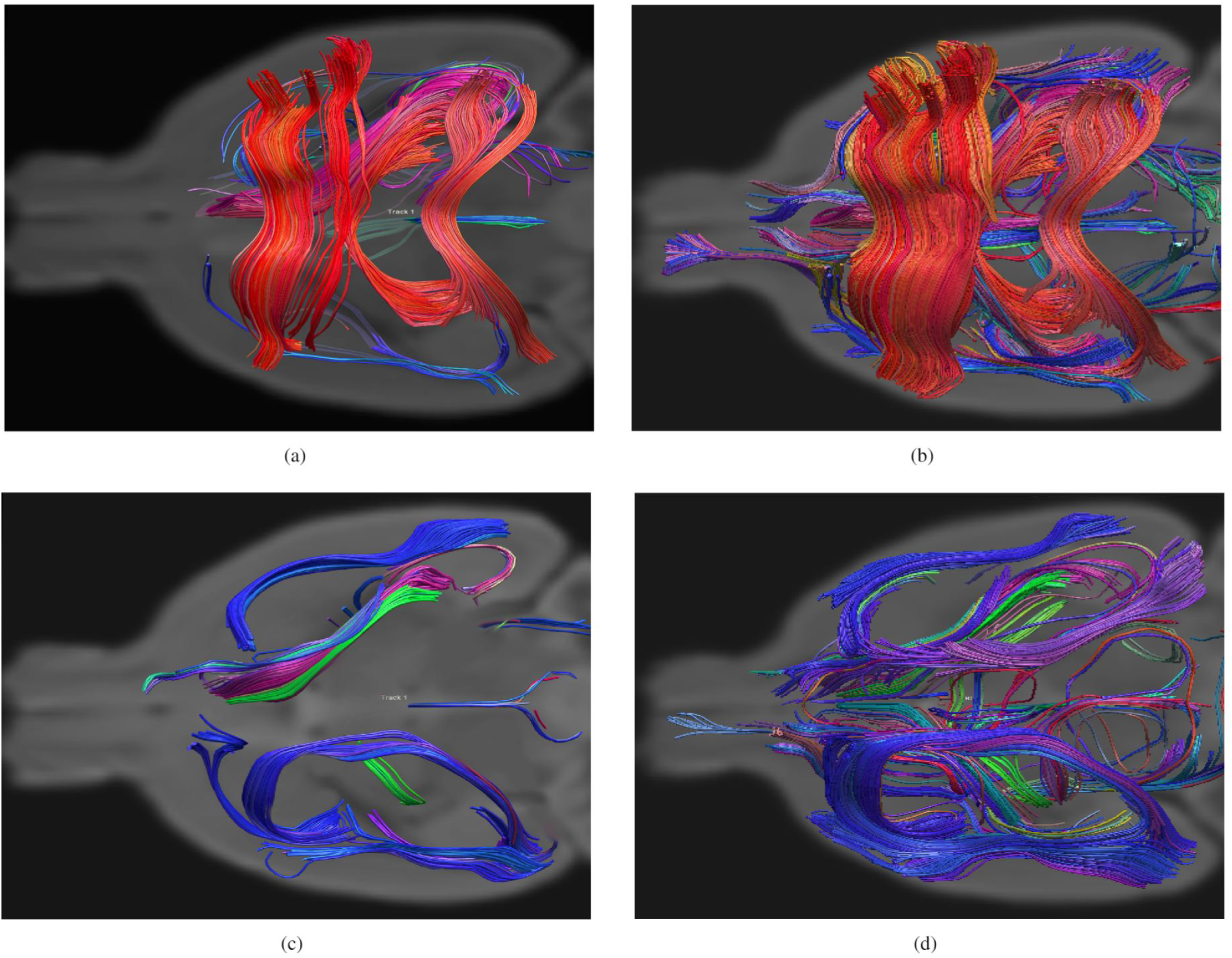
Graphical representation of the most significant features charactering the structural connectome of the two populations: the axial views of a randomly selected subject from the C57BL/6J control population (a) using our algorithm (***α*** = 110) and (b) the NBS algorithm using as a threshold p-value 0.05. While (c) and (d) are the axial views and of a randomly selected subject from the BTBR population respectively for our algorithm and NBS. As expected BTBR mice show a lack of corpus callosum and hippocampal commissure and an increased intra-hemispheric ipsilateral connectivity. Performing the same experiments by using the SVM framework, similar results if using NBS where obtained. The depicted discriminant features detected by the proposed algorithm can be increased varying the ***α*** parameter according to the user taste.

Similar results were obtained with the structural dataset on Alzheimer’s disease. Altered brain connectivity both at the microstructural and macrocircuitry levels has been described in this disorder due to the amyloid plaques^53^. As depicted in Figure 5, we found widespread temporal and para-hippocampal connectivity differences in patients with Alzheimer’s disease compared to healthy elderly subjects. This is inline with several studies about functional connectivity which highlighted decreased functional connectivity between the temporal gyrus and neighbouring regions^34,54^. Pathways between the hippocampus, the parahippocampal gyrus and neocortical regions are considered to be the first affected in Alzheimer’s disease patients^55,56^. Neurodegeneration and loss of connectivity between frontal areas, the insula and within the areas of the frontal gyrus has also been documented^57^. The found connection between the cuneal cortex left and the frontal opercolum right were possible through fibers bifurcating from the corpus callous. Studies have shown that the functional activities at the cuneal cortex is reduced with the progress of Alzheimer^58^, this can explain the visual field defects seen in some patients^59^.

**Figure 5.**
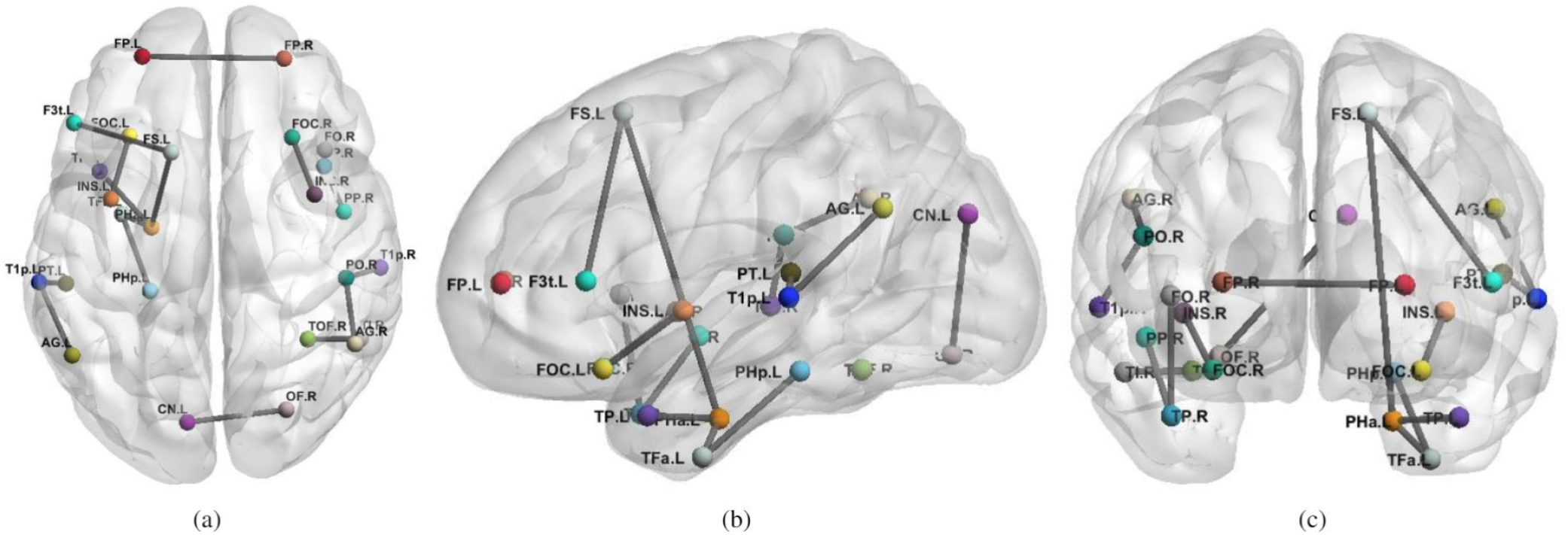
Structural connections differentiating patients with Alzheimer from elderly controls with MLA (our algorithm) using ***α*** = 33. From left to right, axial (a), sagittal (b), coronal (c) views of the brain indicate significant connections not set to zero by the algorithm. Each line represents a specific structural connection. The acronymous are the same as reported in Table 1.

Regarding the discriminant connections detected by the MLA algorithm for the ADHD dataset, among the detected areas using α = 10, there were the connections between the Frontal Pole and the Cingulate Gyrus, and the Frontal Pole and Angular Gyrus, which are the main functional hubs of the default mode network (DMN). The DMN is known to be altered in ADHD subjects^60,61^. As it has been hypothesized that ADHD subjects may have diminished ability to inhibit the default processing of the DMN^62^. The other detected connections could be explained as dorsal medial and medial temporal systems still related to the DMN^63^.

The connectivity differences for the human dataset, shown in Figure 5 and 6, and reported in Table 1 and 2 were those detected by MLA, some of which were not discovered by NBS and SVM. This shows that different approaches - one based on sparsity and one based on family-wise error rate can produce similar results, though the proposed method can find more features than the NBS or other univariate approaches. NBS and MLA select discriminative features in different ways. NBS performs univariate t-tests among the features while MLA performs a sparse multivariate regression. It is acknowledged that the methods achieve different objectives. However, it has been shown that sparse model jointly with a stability selection criteria can lead to more robust results than models based on false discovery rate or family-wise error-rate^64^. The advantage of MLA resides on the possibility to use it in a exploratory framework, where it provides insights (detected connections) to neuroscientists for a deeper investigation.

**Table 1.**
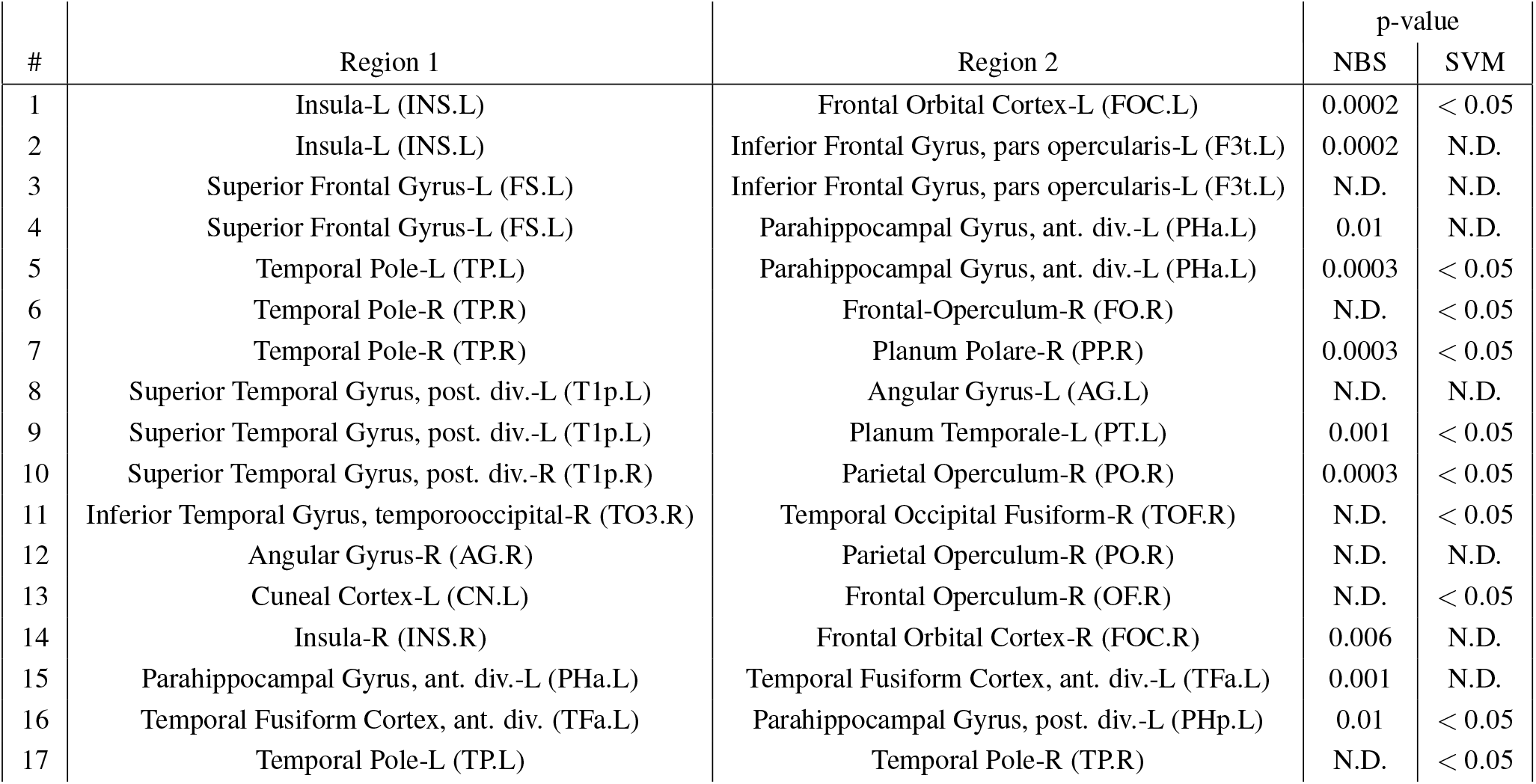
Structural connections differentiating patients with Alzheimer from normal elderly individuals detected by MLA. Pairs of source and target regions and p-values of the univariate t-test computed on NBS^26^ and SVM weights using the t-test threshold corresponding to p-values < 0.05^20,21^ are reported. “Not detected” (N.D.) means not significant difference between the two areas.

**Table 2.**
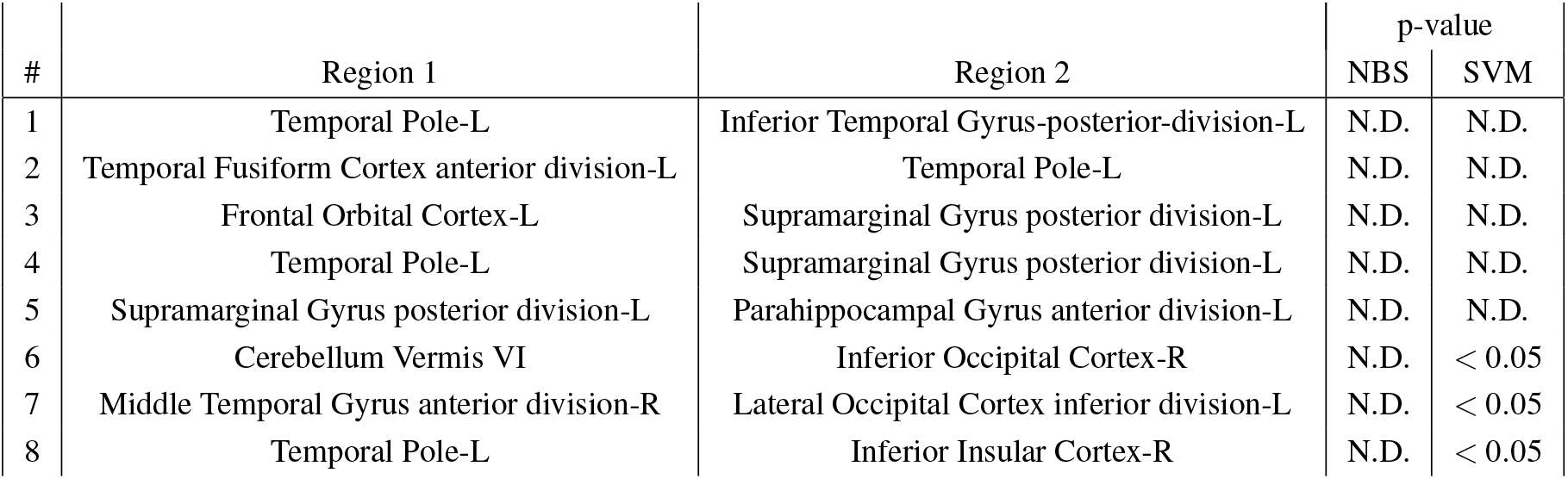
Functional connections differentiating patients with ADHD from TD individuals. Pairs of source and target regions and p-values of the univariate t-test computed on NBS^26^ and SVM weights^20,21^ are reported. “Not detected” (N.D.) means not significant difference between the two areas. ADHD= Attention-Deficit/Hyperactivity Disorder, TD= Typically developed. For this experiment, no statistically significant features were obtained by the NBS and SVM-based algorithm using the t-test threshold corresponding to p-value < 0.05^20,21^.

| # | Region 1 | Region 2 | p-value |  |
| --- | --- | --- | --- | --- |
|  |  |  | NBS | SVM |
| 1 | Temporal Pole-L | Inferior Temporal Gyrus-posterior-division-L | N.D. | N.D. |
| 2 | Temporal Fusiform Cortex anterior division-L | Temporal Pole-L | N.D. | N.D. |
| 3 | Frontal Orbital Cortex-L | Supramarginal Gyrus posterior division-L | N.D. | N.D. |
| 4 | Temporal Pole-L | Supramarginal Gyrus posterior division-L | N.D. | N.D. |
| 5 | Supramarginal Gyrus posterior division-L | Parahippocampal Gyrus anterior division-L | N.D. | N.D. |
| 6 | Cerebellum Vermis VI | Inferior Occipital Cortex-R | N.D. | $< 0.05$ |
| 7 | Middle Temporal Gyrus anterior division-R | Lateral Occipital Cortex inferior division-L | N.D. | $< 0.05$ |
| 8 | Temporal Pole-L | Inferior Insular Cortex-R | N.D. | $< 0.05$ |

**Figure 6.**
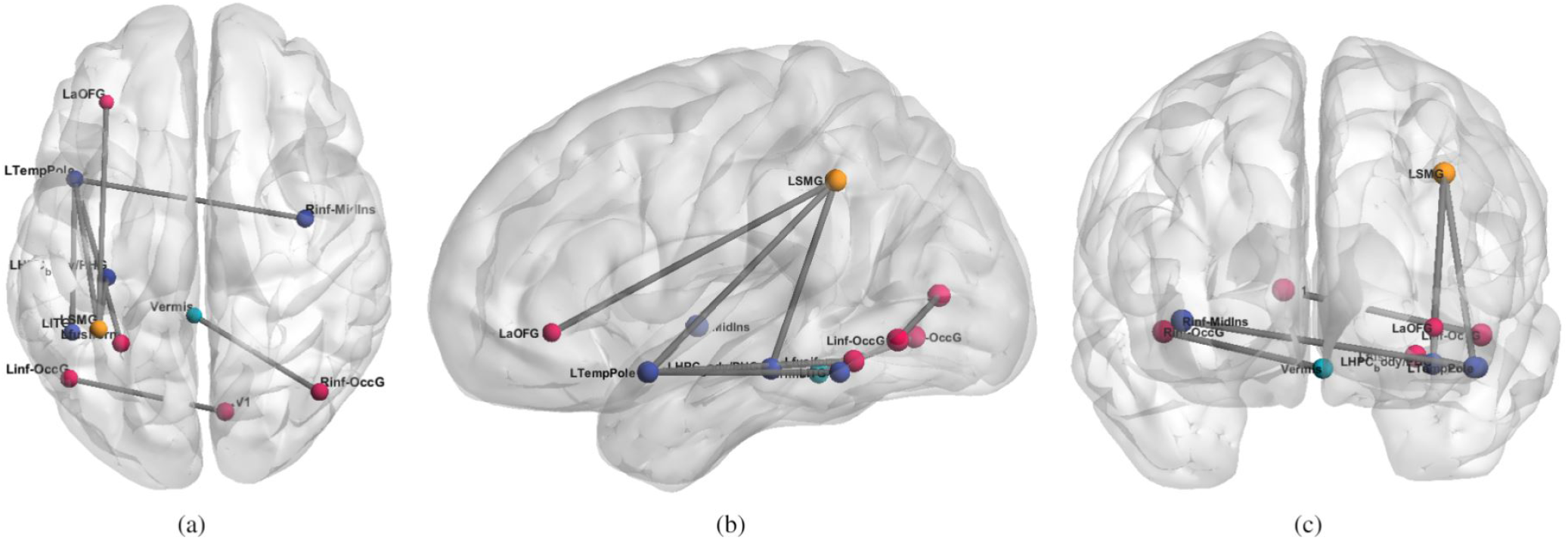
Functional connections differentiating ADHD from TD subjects obtained by using the proposed method (MLA) using ***α*** = 10. From left to right, axial (a), sagittal (b), coronal (c) views of the brain indicate significant connections not set to zero by the algorithm. Each line represents a specific functional connection. For details on the statistics and name abbreviations, see Table 2.

## Conclusions

In this manuscript, a fully automated method to characterise brain connectivity in case-control studies was reported. The method based on a sparse learning classification, has been tested on structural and functional connectivity data. The approach is able to identify brain areas of interests that can be further analysed with standard seed based approaches or through histological white matter validation.

The algorithm successfully highlighted some known structural white matter differences in acallosal mice, and identified previously reported alterations of structural and functional connections in human Alzheimer’s and ADHD patients. The developed software is freely distributed as a Matlab toolbox at the url *https://github.com/alecrimi/multi-link*. Our approach can help highlighting differences in connectivity generating hypotheses that can complement univariate techniques.

## Methods and Data

This section first describes the two types of data used to test the proposed method: a mice dataset with high dimensionality, and two publicly available human datasets. Afterwards the pre-processing and the proposed computational model for discriminating patterns in whole-brain analysis are described.

### Data

#### Mouse Structural Connectivity Data

The mice cohort was composed of two groups of 22-26 weeks old male subjects (n=16): BTBR T+tf/J mice (n=8) which share analogies to all diagnostic symptoms of autism and characteristic functional and structural features of the brain^47–49^, and C57BL/6J mice (n=8) which are characterised by normal sociability and represent the control group. Figure 1 depicts an example of the expected difference between the BTBR and C57BL/6J mice groups. In particular, BTBR mice lack the corpus callosum differently from the C57BL/6J mice.

The animal preparation protocol has been already described^45,49^. Briefly, brains were imaged inside intact skulls to avoid post-extraction deformations. Ex-vivo high-resolution DTI and T2-weighted images were acquired on paraformaldehyde-fixed specimens with a 7 Tesla Bruker Pharmascan MRI scanner (Billerica, MA, USA). T2-weighted MR anatomical images were acquired using a RARE sequence with the following imaging parameters: TR/TE = 550/33 ms, RARE factor = 8, echo spacing 11 ms, and a voxel size of 90 *μm* isotropic. DTI volumes were acquired using 4 scans at b0 and 81 scans with different gradient directions (b=1262 s/mm^2^), with resolution 130 × 130 *μm*^2^, using a 4-shot EPI sequence with TR/TE = 5500/26 ms. Anatomical and DTI sequences were acquired sequentially at the same centre with the same scanner. This dataset is freely distributed^65^. This dataset is used to show that the algorithm is able to identify difference between the groups which are expected to be found as a proof of concept.

#### Human Structural Connectivity Alzheimer Data

The human experiments have been performed on the Alzheimer’s Disease Neuroimaging Initiative (ADNI) dataset publicly available^66^. Only baseline scans were used to avoid confounding factors as advanced brain atrophy and treatment in the Alzheimer patients. This cohort comprised 51 Alzheimer’s Disease patients (age: 76.5 ± 7.4 years), and 49 normal elderly subjects (77.0±5.1) matched by age. The used data were DTI, and T1-Weighted obtained by using a GE Signa scanner 3T (General Electric, Milwaukee, WI, USA). The T1-weighted scans were acquired at with voxel size =1.2 × 1.0 × 1.0 *mm*^3^ TR = 6.984 ms; TE = 2.848 ms; flip angle=11°). DTI were acquired at voxel size =1.4 × 1.4 × 2.7 *mm*^3^, scan time = 9 min, and 46 volumes (5 T2-weighted images with no diffusion sensitization b0 and 41 diffusion-weighted images b=1000 *s/mm*^2^). For each subject, DTI and T1 have been acquired and co-registered.

#### Human Functional Connectivity ADHD Data

Functional connectivity was also investigated on a larger resting-state fMRI dataset comprising ADHD and TD subjects^67^. In particular, we used the publicly available New York University Child Center dataset^68^, which is the main cohort of this study. The dataset comprised 95 ADHD subjects (67 male and 28 female, mean age 11.4 ± 2.7) which were either inattentive, or hyperactive or both, and 92 healthy TD (45 male and 47 female, mean age 12.4 ± 3.1) which represents the control group.

The fMRI volumes were acquired with a Siemens Allegra 3T, with TR/TE 2000/15 ms and voxel size 3 × 3 × 3*mm*^3^. This dataset is used as a real case study where NBS and other univariate approaches are not useful in proving local connectivity differences.

### Methods

#### Mouse Dataset Processing and Encoding

Deterministic tractography was performed on the DTI volumes by using the software tool DiPy^69^ after eddy current corrections, by using the Fiber Assignment by Continuous Tracking (FACT) algorithm^70^. Fibres were reconstructed in the original volumes following the 2nd-order Runge-Kutta integration scheme^71^ starting from the centre of each voxel and following the main direction of the tensor. The tracking was stopped when the fibre made a sharp turn (> 35°) or entered a voxel with fractional anisotropy (FA) < 0.15.

To allow inter-subject comparisons, registration matrices to a common space were computed for each subject by using affine transformation (12 degrees of freedom). The obtained registration matrices were then applied to the endpoints of each fibre. This allowed the tractography algorithm to work on the original volume space without warping the tensors.

To enable a purely data-driven inter-group comparisons without the use of anatomical priors, the brain volumes were split into 3D cubes of size 1 × 1 × 1*mm*^3^, without considering any atlas. Each cube was a node in the graph and the connectivity matrix was built counting the fibres starting and ending into two distinct cube elements of the grid, avoiding the inclusion of u-fibers. This resulted in defining 42,704 edges.

The advantage of this approach was that the result of the proposed analysis method was nearly independent from the size and the type of parcellation. Indeed, not considering the anatomy nor the physiology of the brain might result in bundles of fibers split into “sub-bundles” connecting adjacent cubes. However, if there is a difference between the two groups it is retrieved for all sub-bundles, hence the overall bundles are then reconstructed. Yet the choice of using a fine grid or an atlas is arbitrary.

#### Alzheimer Dataset Processing and Encoding

Tractographies for all subjects have been generated processing the DTI data with a deterministic Euler approach of DiPy^69^, stemming from 2,000,000 seed-points and stopping in case of FA smaller than 0.25. Tracts shorter than 3cm were discarded during the connectome construction. Structural connectivity matrices were constructed by counting the number of fibers connecting two regions of interest (ROIs) of the registered Harvard-Oxford atlas^72^. This atlas defines 96 ROIs, it is freely available with several brain imaging analysis platforms, and it has been used in several structural studies including Alzheimer’s Disease^73^. The Harvard-Oxford atlas is a probabilistic atlas. However, we used the version where each voxel is associated to the ROI with highest probability. The choice of different algorithms used for the tractography with the human and murine data is related to the fact that the data are obtained with different types of scanner: a small animal device and a common clinical scanner.

#### ADHD Dataset Preprocessing and Encoding

This dataset has been pre-processed^60^, and the final connectivity matrices are publicly available^68^. In brief, resting-state fMRI data were preprocessed following these steps: Removal of first 4 EPI volumes, slice timing correction, motion correction, and then applying the regressors for WM, CSF, motion time courses and a low order polynomial detrending. A band-pass filter of 0.009 < *f* < 0.08 Hz was also applied. Lastly, the data were blurred using a 6-mm Full Width at Half Maximum Gaussian filter. The functional region of interests were obtained using the Craddock parcellation^74^ for 200 areas. Those preprocessing steps have been carried out according to the Athena pipeline^75^ which is based on a combination of command from AFNI and FSL.

### Parameter Tuning

While the number of discriminative connections selected by our type of model is tuned by the choice of ***η***, we noticed that the algorithm was satisfactorily discriminating the two classes on a wide range of ***η*** values. In this work we were mostly interested on discriminant features rather than finding an optimal classification. However, the results are shown using the values which allow better accuracy estimated in a nested cross-validation manner to produce a jackknife-like classification. In practice a nested leave-one-out procedure was employed. For 1000 iterations, a sample was removed from the training dataset and used as test-set to find the optimal value within a range, then the performances were evaluated on another sample also removed prior the optimization from the training set and used as validation. In presence of a plateau of identical optimal values, the value generating less connections was taken. The range of values has been previously identified empirically. Namely, several values have been tried looking for those showing meaningful connectivity.

The optimal parameter ***γ*** was investigated in a grid search for all experiments. It was noted that, conversely to the parameters of the other models, the value of this parameter is not critical for the classification. Nevertheless, values larger than 0.03 were producing slightly better results than smaller. Lastly, the parameter ***η*** can also be reformulated as the desired number of variables selected by the model. In the following we will refer to this number as ***α*** instead of ***η***. We address the reader to Zou et al.^41^ for a description of the relation between ***η*** and ***α***, and further details on the algorithm are given in the Appendix.

Although this model can be very powerful in determining small and good subset of features allowing to linearly discriminate the classes, it suffers from a stability problem^76^, i.e., small changes in the data could change the result of a single run. To cope with this stability issue, in order to improve the robustness of SDA, we have introduced a second stage exploiting the ensemble of low-stability algorithms to produce a more stable feature selection. In practice, we perform further feature selection to ensure that the features are stable across subjects.

In the specific case, the SDA classifier was trained with a nested leave-one-out approach. This ended in an ensemble of models each one with a subset of “relevant” features (connections), selected so to maximize the discrimination between the two groups. Then we refined this ensemble of models by occurrence validation, where only features which were frequently selected during cross-validation were retained, i.e., features occurring in less than a pre-defined percentage of runs were discarded.

In all the experiments reported in the paper this threshold was determined as half the number of subjects in the corresponding dataset. In this way, we ensure stability selection of the features^28^. The choice of using half the number of subjects as threshold for sampling features has been already used^64,77^ as a trade-off between considering all features (no restriction) and considering only features which occurs in all samples (over-restrictive). The impact of varying the threshold has already been investigated^64^, showing that a small number of selected features (obtained by thresholding) guarantees a small number of false positives. Nevertheless, the focus of the paper is on obtaining an optimal true positive rate more than a low number of false positives.

## Author contributions statement

A.C., L.G., V.M. and D.S. conceived the experiments, A.C. and L.G. conducted the experiments, A.G. acquired the murine data, A.C., L.G., F.S. and D.S. analyzed the results. All authors reviewed the manuscript.

## Appendix

### Sparse Discriminant Analysis

The general formulation of *ℓ*_1_ regularization or *lasso* is used in regression frameworks to minimize the problem *min_β_*{||**y** − **X*β***||^2^ + ***η***||***β***||_1_}, where **X** is a data matrix, **y** is the output vector, and ***β*** is the regressor vector. Similarly, the elastic net is given as *min_β_*{||**y − X*β***||^2^ + ***η***||***β***||_1_ + ***γ***||***β***||_2_}. In these equations ***η*** and ***γ*** are tuning parameters which are used to yield sparse coefficient vector estimation^41^. The parameter ***η*** can also be reformulated as the number of desired variables which are left in the model, when used in this context we refer to it as ***α***^41^.

There are several extension to the linear discriminant analysis^78^ which comprises Lasso and elastic net^40^. Our experiments are based on the formulation proposed by Clemmensen et al^30^. More specifically, given the matrix data **X** with *n* p-dimensional observations for K=2 classes each of them defined as **x_i_**, with ***μ***_**k**_ representing the mean for a specific class *k*, it is possible to define the within-class covariance matrix common to all classes as 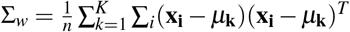, and the between-class covariance matrix 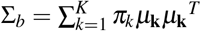, where ***π***_*k*_ is the prior probability for each class to belong to the class *k*. The prior probability is generally given by the size of respective classes.

A Fischer discriminant analysis can classify to which class a sample belongs by using discriminant vectors whose directions ***β***_**k**_ maximize

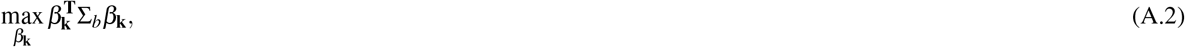

subject to 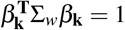 and 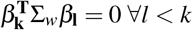.

Very often, as in our case, the previous maximization process is ill-posed, as the matrix Σ_***w***_ might not be full rank as the number of features is far larger than the number of available samples. A possible solution, proposed by Witten et al.^40^, is given by using the Lasso or elastic net regularization as

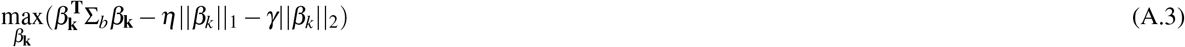

subject to 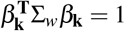 and 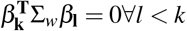. Alternatively we used the minimization formulation of Clemmensen et al.^30^, where the pair given by ***β***_**k**_ and the vector of scores ***θ***_**k**_ solves the problem

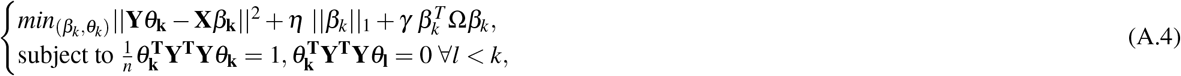

where Ω is an arbitrary positive matrix, ***η*** and ***γ*** are non negative tuning parameters, and **Y** is a *n* × ***K*** matrix of dummy variables for the K classes. This formulation of LDA as a regression problem introduces sparsity, and allows its use when the number of features is very large compared to the number of available samples.

